# Exploring Contextual Interference in Implicit and Explicit Motor Learning

**DOI:** 10.1101/644211

**Authors:** Kristy V. Dang, Darius E. Parvin, Richard B. Ivry

## Abstract

The classic advice given to anyone learning a new skill is “practice makes perfect.” While this provides a good general rule to follow, it lacks any detail on what form of practice will efficiently maximize learning. So when faced with the task of acquiring multiple skills, what is the optimal way to learn? Would it be more beneficial to master each skill separately or learn them all at once in an interleaved fashion? A concept known as contextual interference suggests that using a random practice schedule leads to better retention than a blocked one. There are some motor learning studies that are consistent with this hypothesis and some that are not. In order to explore these conflicting results, we applied contextual interference to a simple reaching task that could allow us to observe its effects to various components of motor learning. We had participants learn three different visuomotor rotations and manipulated interference by placing them in groups characterized by how training targets are ordered (blocked vs. random). Using reaction time and hand angle as our measures of performance, we found that participants who experienced a random practice schedule had significant improvements in their ability to retain information, which was manifest as higher levels of implicit adaptation and faster reaction times. However, this did not necessarily mean the information was executed accurately since hand angles did not differ between groups. These findings suggest contextual interference will be most advantageous in situations that require fast explicit recall of a motor plan to use rather than those that emphasize accuracy.

## Introduction

Teaching a new skill could be tricky because students may require tailored lesson plans. However, there is considerable interest in finding methods that optimize learning despite these individual differences. This would allow teachers and coaches to utilize a training framework that could then be built upon depending on the situation. Using a series of verbal tests, Battig (1972) laid out a learning theory called contextual interference. He argued that retention can be improved if there is a high amount of interference during the initial training period. He believed that experiencing disruptions makes learning less context-dependent, forcing the individual to be more highly engaged in the task.

Shea and Morgan (1979) were the first to apply the idea of contextual interference to a motor learning task. They utilized a unique set-up of six wooden barriers (2×3) that participants had to quickly knock down using one of three pre-determined sequences. Each one was associated with a colored light that would turn on and indicate which order of movements to do. For example, a blue light meant that participants would have to sequentially knock down the right rear, left middle, and right front barriers, while a red light signaled right front, left middle, and right rear. They introduced contextual interference into their experiment by differing the participants’ training schedule. One group (Blocked) experienced low interference by learning each sequence in separate blocks of training, while the other group (Random) experienced high interference by learning all three sequences simultaneously in a random order. The Blocked group initially outperformed the Random group during training. This was expected since they only had to remember one sequence at a time. By the end of the practice session, there was little difference between the groups’ performances, showing that all participants fully mastered the task. After a delay, participants were retested to assess how much they remembered. Overall, the Random group had higher levels of retention despite their poor performance during training, consistent with the hypothesis that interference can be beneficial for long-term learning. This showed that the idea of contextual interference could be used to enhance motor skill learning.

This finding spurred many follow-up studies with mixed results. There have been numerous controlled, laboratory experiments using a multi-segment movement task similar to that used by Shea and Morgan. These studies have consistently reproduced the contextual interference effect (Lee & Magill, 1983; Gabriele et al., 1987; 1989; Shea and Zimny, 1988; Magill & Hall, 1990). There have also been real-world, non-laboratory experiments that try to extrapolate the benefits of interference to more common motor activities. However, their results are not as consistent. Goode and Magill (1986) found that random participants outperformed during retention in a study on badminton serves, but Landin and Herbert (1997) could not replicate these results in an experiment that instead used basketball free-throws from various locations on the court. Similarly, experiments using various volleyball skills showed a lack of retention benefits for the high interference group (French et al., 1990; Bortoli et al., 1992). In fact, there seem to be few studies that show the contextual interference effect in a real-world setting. This motivates a more nuanced question: Which motor learning processes benefit from practice schedules that are highly variable?

To explore this idea, we used a contextual interference design in a basic sensorimotor learning task that allows us to measure different features of learning. Participants learned to compensate imposed perturbations of their visual feedback as they reached to different targets. These simple reaching tasks have been shown to involve at least two distinct motor learning processes that operate in parallel of each other. (Mazzoni & Krakaeur, 2006). (1) An implicit process, dependent on the cerebellum, is important for unconsciously recalibrating the body’s movements (Tseng et al., 2007; Taylor et al., 2010; Taylor & Ivry, 2014). (2) Explicit learning, on the other hand, focuses more on the conscious and deliberate use of strategies, which has been shown to depend more on the prefrontal cortex (Anguera et al., 2010; Taylor & Ivry, 2014). Together, these two processes generate more precise movements by explicitly selecting a movement and then implicitly fine-tuning the execution of the movement. In the present study, we use this basic reaching task (also known as a visuomotor rotation experiment) in order to break down these motor learning components and understand if and how contextual interference differently affects them.

We had participants learn to compensate for three different rotations. Each rotation was associated with a target that appeared at a unique location in space. We manipulated the degree of interference by varying the training schedule. For the Random condition, the three targets (and thus rotations) were presented in an interleaved, random fashion. For the Blocked condition, the three targets were presented in a sequential manner. If contextual interference was operative in this learning task, we would expect to find the following: 1) Those experiencing a random practice schedule should perform poorly during training but 2) would outperform the blocked participants at retest. According to Battig’s initial hypothesis that the advantages of contextual interference are due to a greater conscious engagement of mental processes, we would expect to see most of this interference effect in the explicit learning portion of the task.

## Methods

### Participants

A total of 72 undergraduate students (53 females, age = 18-25 years old) were recruited from the University of California, Berkeley. All were neurologically healthy with normal or corrected vision and verified to be right-handed through the Edinburgh Handedness Inventory (Oldfield, 1971). Participants were financially compensated for their participation. The experimental procedure was approved by the Institutional Review Board of UC Berkeley.

### Experimental apparatus

Participants held and moved a stylus-embedded air hockey paddle on top of a digitizing tablet (49.3 cm by 32.7 cm, Intuos 4 XL; Wacom, Vancouver, WA) that continuously recorded the position of the stylus at 200 Hz. An LCD screen (53.2 cm by 30 cm, ASUS) mounted above the tablet served to display the visual stimuli as well as occlude the participant’s hand from view. The experimental task was written in Matlab and displayed using Psychtoolbox extensions (Pelli, 1997).

### Experimental design

#### Visuomotor Reaching task

The participant was instructed to make horizontal reaching movements from a starting location (white circle, 6 mm) to targets (blue dots, 6 mm) that appeared at various positions on the screen. A white cursor (3.5 mm) provided visual feedback of their hand position. Targets appeared in one of three locations (30°, 150°, or 270°), and their order of appearance depended on experimental conditions described below (Figure 1A).

**Figure 1:**
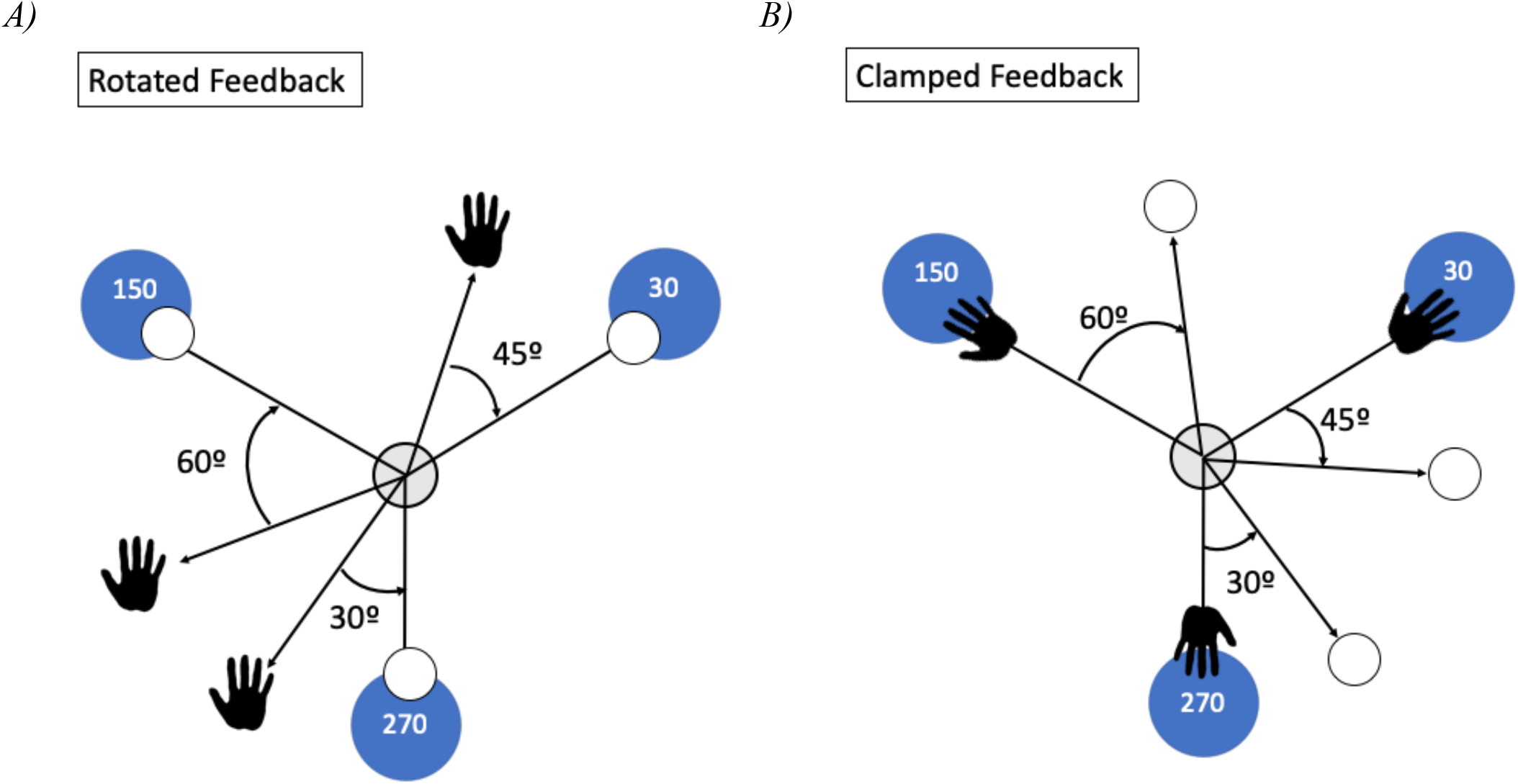
Example Rotation Condition. **A)** Rotated feedback: in Experiment 1, the cursor was rotated relative to the participants’ hand position. There were 6 possible rotation conditions, one of which is shown here: 45° counterclockwise, 60° counterclockwise, and 30° clockwise for the 30°, 150°, and 270° targets respectively. The hand indicates where participants would need to reach in order to compensate for the rotation and land the cursor onto the target. **B)** Clamped feedback: in Experiment 2, the cursor was on a fixed path irrespective of the participants’ hand position. The rotation conditions were identical to the ones in Experiment 1. Participants were told to ignore the rotation on the cursor and move their hand directly to the target. The hand indicates where participants should be reaching despite the rotated feedback.

Participants would begin every trial by moving their hand into the starting position. When their hand was 2 cm from the center, a white search ring would appear with a radius that was correlated to the distance of the participant’s hand to the desired starting position. The ring would become smaller the closer the participant got to the center, which helped to guide their movement in the right direction. Once participants were 1 cm away, the ring would be replaced by a white cursor that directly corresponded to the position of the hand. Participants would have to hold the cursor within the start location for 500 ms before a single target would appear and they could then begin their reach. These movements were instructed to be quick and straight through the target. Movement onset was defined as the time when movement amplitude reached 1 cm from the center of the start position, and movement time (MT) was defined as the interval between movement onset and the time at which the hand reached the radial target distance (8 cm). Auditory sounds were played to encourage consistent MTs: a pleasing knock for correct speeds, the words “too slow” if the reach exceeded 300 ms, or the words “too fast” if the reach was under 100 ms.

For trials with visual feedback, the cursor was visible throughout the reach until the hand exceeded the radial target distance. At this point, the endpoint position of the cursor was frozen for an additional 1 s to allow participants to receive information about their performance accuracy. All feedback on the screen would then disappear, and participants would move their hand back to the start position to begin the next trial.

#### Rotation conditions

Following a series of baseline trials, a visuomotor rotation was applied so that the location of the cursor no longer corresponded with the participants’ hand position. In Experiment 1, the cursor was rotated at a fixed angle from the actual hand position (Figure 1A). The participants’ goal was still to hit the target with the cursor, so they had to re-strategize the direction in which they reached their hand. In Experiment 2, the cursor had a task-irrelevant clamped feedback that was rotated away from the target and followed a fixed path independent of hand position (Figure 1B). Participants were instructed to ignore the feedback and just “reach directly to the target with their hand.”

To reduce the ability of participants to generalize their reaching strategy of one target to another, we used rotations that varied in magnitude and direction for each target. Everyone was given one of 6 possible rotation sets that included three rotations, one for each of the three targets. For example, if a participant was given the set [45°, 60°, −30°] that meant that the cursor would be rotated 45° counterclockwise for the 30° target, 60° counterclockwise for the 150° target, and 30° clockwise for the 270° target. Since all of the rotations were different, the participant had to develop an individual reaching strategy for each target. The possible rotation sets for the 30°, 150°, and 270° targets respectively were: [45°, 60°, −30°]; [30°, −45°, −60°]; [−30°, 60°, 45°]; [−60°, 30°, −45°]; [60°, 45°, −30°]; [−45°, −60°, 30°] (positive indicates counterclockwise direction, while negative indicates clockwise; first rotation combination shown in Figure 1).

#### Experiment 1

The goal of Experiment 1 was to directly assess the effects of a random versus blocked practice schedule on learning in a visuomotor reaching task. Participants were randomly assigned to either the Random (n=18) or Blocked (n=18) group. The difference between the groups was in how the targets were ordered during the training period (Figure 2A).

**Figure 2:**
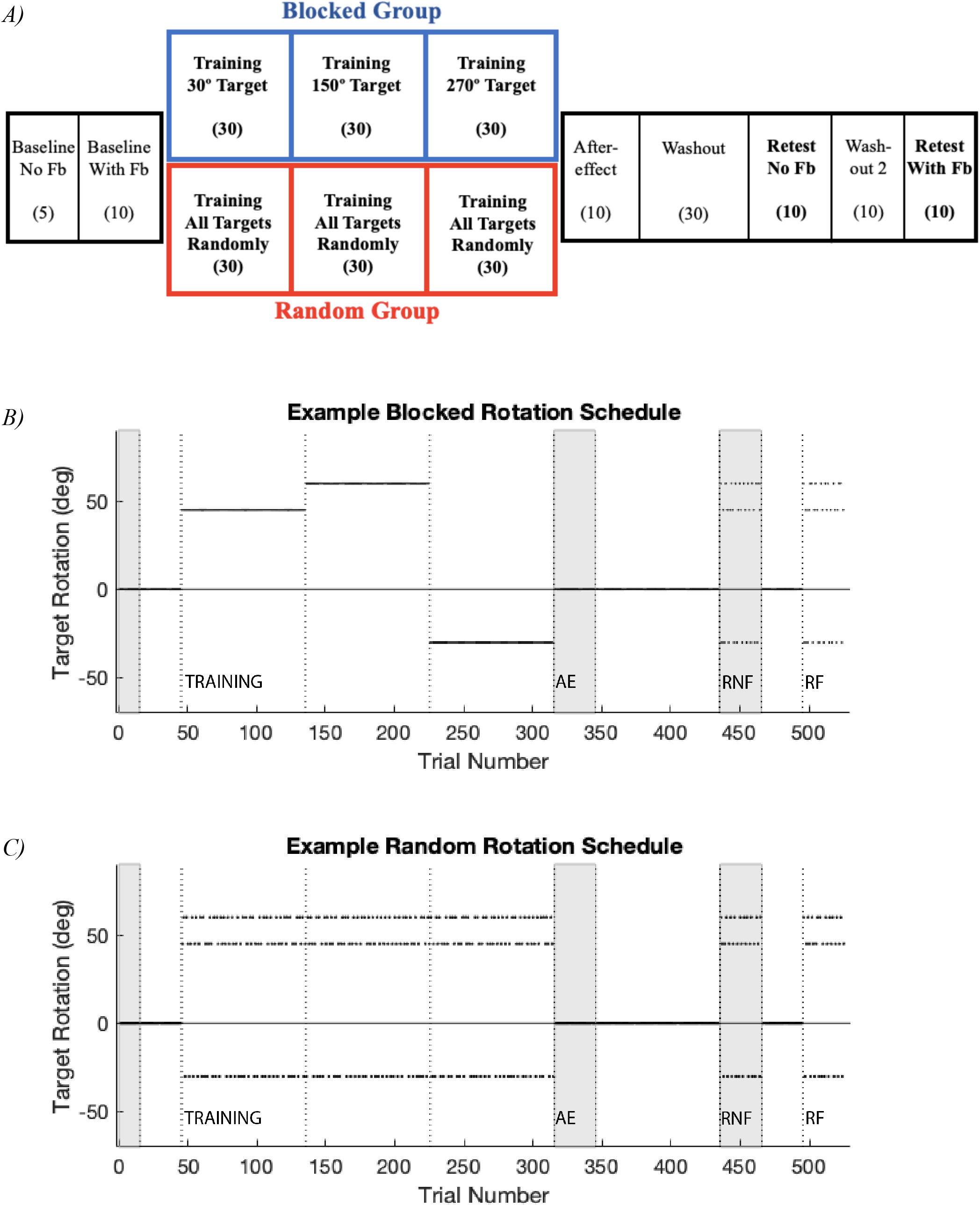
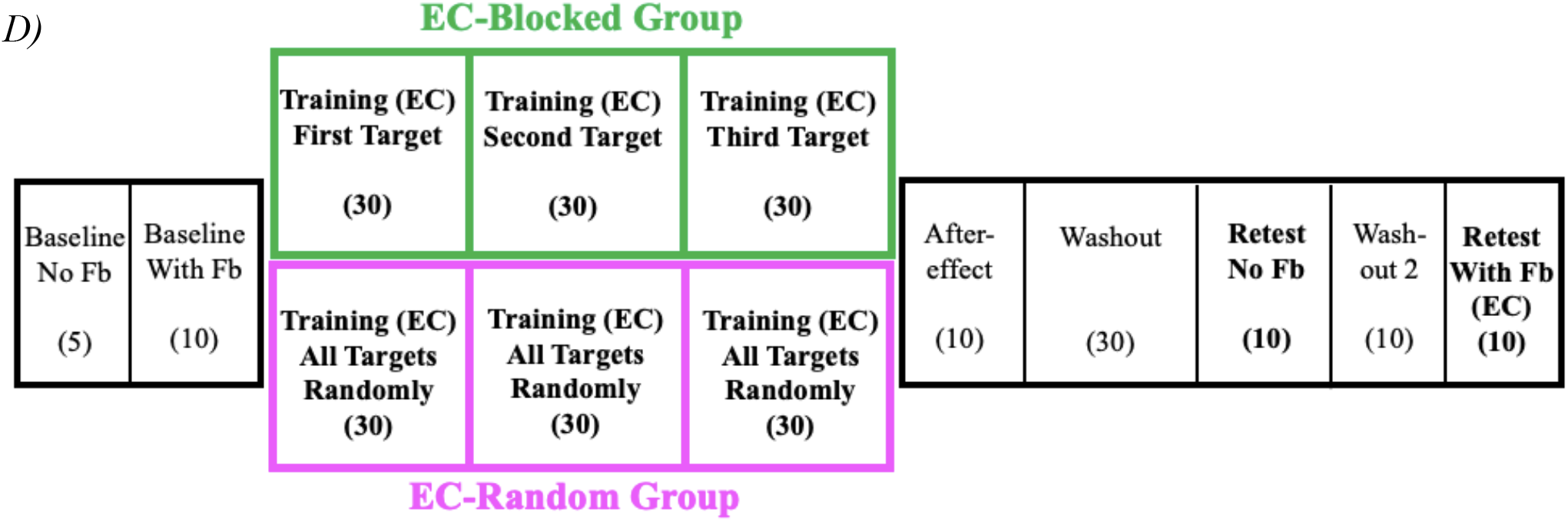
Task design. **A)** Participants in Experiment 1 placed in either the Blocked or Random group. All participants experienced the same pre- and post-training schedule. The number in parentheses indicate the number of cycles in each block. **B/C)** Example Blocked and Random rotation schedules: Training, Aftereffect (AE), Retest No Feedback (RNF), and Retest With Feedback (RF) blocks labeled. No feedback blocks shaded. Same rotation condition as shown in Figure 1 **D)** Participants in Experiment 2 were similarly placed into two groups: EC-Blocked and EC-Random.

Both groups started with 5 cycles of baseline trials (each cycle is composed of one reach to each of the three targets) with no visual feedback and 10 cycles of baseline trials with veridical feedback. For the Training block, participants were informed of the target order structure and the multiple rotations that would be imposed on the cursor for each target. They were instructed to find the appropriate strategy required to hit the targets with the cursor and to remember any successful strategies since they would have to reuse them later in the experiment. For this block, the Blocked group practiced with one target for 90 trials before moving on to the next one (Figure 2B), while the Random group practiced all targets in an interleaved manner, with a break provided every 90 trials (Figure 2C). The order of the targets for the Blocked group was the same for every participant: 30°, then 150°, and finally 270°. This order was not counterbalanced since the order of rotations was already being accounted for. The design of the experiment ensured that all Blocked participants experienced a different sequence of rotations even if the order of targets was kept consistent. When they begin with the 30° target, one participant may start with a 30° rotation, while another one may have a 60° rotation. As a result, this reduced the possibility of order being a confounding variable.

Immediately after training, both groups performed a no-feedback Aftereffect block (10 cycles) in which they were told that all of the rotations were now turned off and that they should reach directly towards the target with their hand. This is where we expected to observe how much implicit adaptation was obtained without further influence from explicit processes. The following block was Washout that had the same structure as the Aftereffect block but with veridical feedback (30 cycles). For the Retest blocks, the experimenter then explained that the same rotations as those that appeared in the Training block would be turned on again, and participants should apply the strategies they used before. The first retest block was Retest No Feedback (10 cycles); participants received no feedback of the cursor or the rotations, which provided an estimate measure of how much explicit knowledge was retained. This was followed by 10 cycles of washout trials and then 10 cycles of retest trials with visual feedback (Retest With Feedback) in order to assess retention.

For every non-training block, including the retest ones, the order of the targets was randomized with no immediate repeats. There was no condition that ordered the targets in blocks during the retest period (in a way that would mirror what training looked like for the Blocked group) because previous studies have shown that no matter how the retest is structured the Random group performs better (Shea & Morgan, 1979).

#### Experiment 2

The purpose of Experiment 2 was to examine the effect of blocked or random practice schedules in a task that isolates implicit adaptation. Previous studies have used a task-irrelevant clamped feedback experiment in order to induce adaptation without the participant’s knowledge (Shmuelof et al., 2012; Vaswani et al., 2015; Morehead et al., 2017; Kim et al., 2018). Experiment 2 was identical to Experiment 1, except for the fact that all of the rotations during the training and retest blocks were clamped at a fixed angle from the target (Figure 2D). Participants were placed in either the EC-Blocked or EC-Random group (n=18 in each; EC for error-clamp) that only differed in the way targets were ordered during the training blocks. During the trials with clamped feedback, the instructions were to completely ignore the feedback of the cursor on the screen and focus on moving directly to the target. Participants were given three demo trials in order to familiarize themselves with the new condition and ensure they understood the directions. Since this was an experiment to look at implicit adaptation, there was no strategy that the participants had to develop or remember for the retest blocks.

### Measuring Performance/Data Analysis

All statistical analyses were performed using Matlab 2016b. In order to assess performance, we looked at two different variables: reaction time and hand angle. (1) Reaction time (RT) was defined as the time between the onset of the target and when the participants’ hand had moved 1 cm from the start position. A lower reaction time indicates better performance because it shows participants were more confident of where to move; they did not have to take the time to figure out what to do before they started reaching. Participants were not explicitly told to minimize their RT. Their only RT requirement was to begin their reaches sometime between the target’s appearance and 5 s afterwards. Otherwise, they would hear an unpleasant buzzer sound to let them know they took too long to start reaching. A 5 s window was large enough to see how differently random and blocked participants used this time to decide on which strategy to use and then act upon it. (2) Hand angle was defined as the angle of endpoint hand position relative to the target.

Trials that had a hand angle that exceeded 90° or a reaction time that was greater than 5 s were deemed outliers and removed from analysis (without affecting the outcome of any statistical test). To analyze any significant differences between the groups, two-tailed t-tests were performed. Additionally, we did a linear mixed-effects (lme) model using the fitlme function in Matlab to assess any fixed effects of practice schedule and cursor rotation on the measures of performance.

## Results

### Experiment 1

For Experiment 1, participants were required to learn three different visuomotor rotations one at a time (Blocked group) or simultaneously (Random group) in order to determine the effects of blocked and random practice schedules on learning. They were instructed of the format of the training schedule and told to determine the appropriate strategies required to hit each target with the cursor. To evaluate contextual interference, participants completed a retest phase in the experiment.

The first measure of performance we looked at was reaction time (RT). Participants with a lower RT were considered to be better at the task since they did not have to spend time to either form or remember the correct hand movement before reaching. We averaged participants’ RT per cycle (3 reaches) across the experiment to see how RT changes (Figure 3A). During the training block, the Blocked group had a significantly lower RT than the Random group, which was expected since they only had to learn and focus on one target and rotation at a time (entire training, all rotations: *t*_(105)_ = 4.8519, *p* < 0.001). This effect persisted to the end of the training period and was present for all three rotations, showing that the unsystematic order of the targets was detrimental to the Random group’s performance (Figure 3B; late training [last 10 reaches to every target]: *t*_(34)_ = 4.0892, *p* < 0.0001 for 30° rotation; *t*_(34)_ = 4.3375, *p* < 0.0001 for 45° rotation; *t*_(34)_ = 4.6336, *p* < 0.0001 for 60° rotation). This difference disappears during the Aftereffect and Washout blocks when the participants were told to reach directly to the target. (Aftereffect: *t*_(105)_ = 1.1811, *p* = 0.2402; Washout: *t*_(105)_ = 1.3662, *p* = 0.1748).

**Figure 3:**
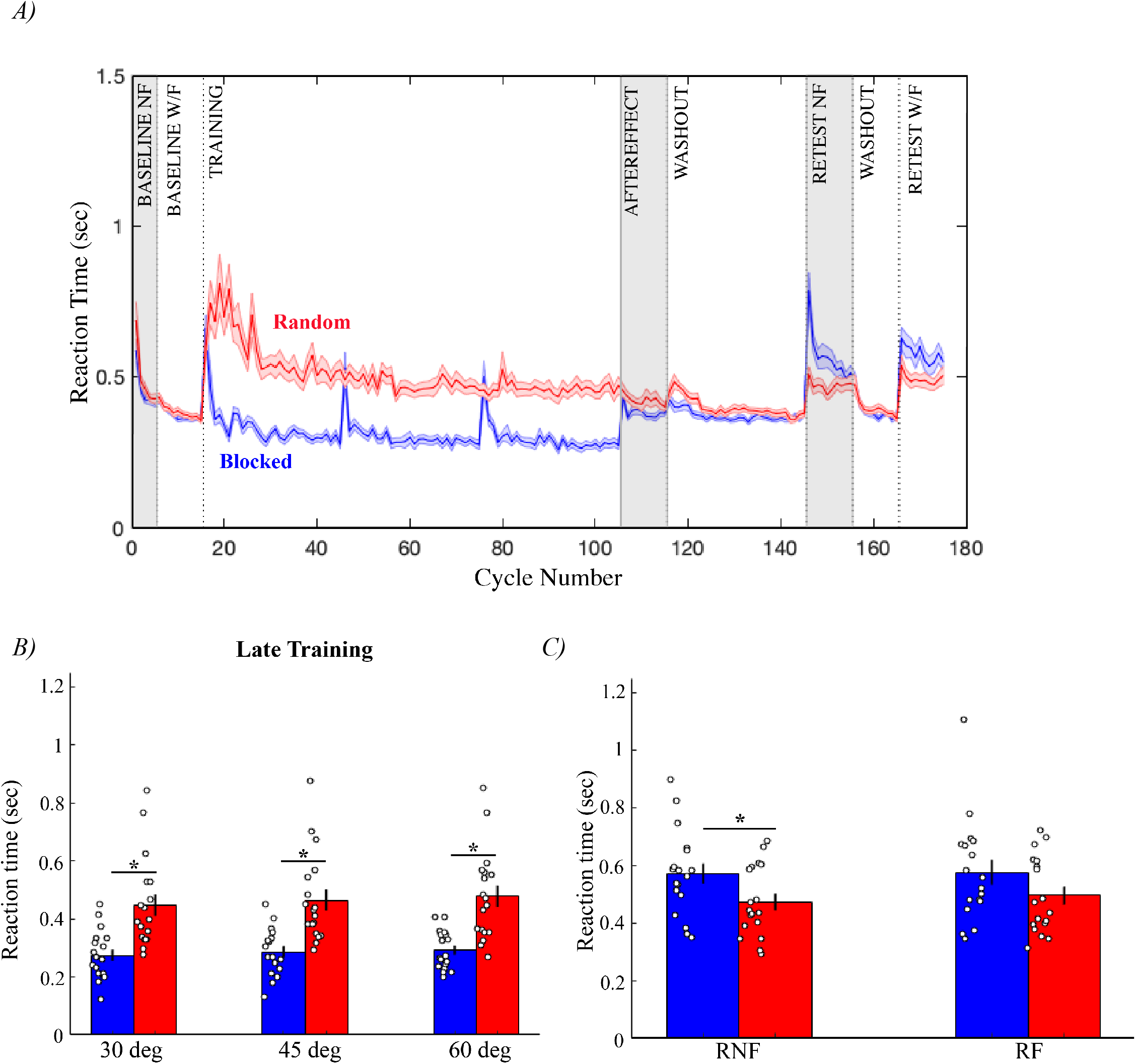
Reaction Time for Experiment 1. **A)** Averaged group RT across Experiment 1. Participant data first averaged by cycle (3 reaches = 1 to every target) before group average is calculated **B)** Late training (last 10 reaches to each target) RTs split by rotation **C)** RNF (Retest No Feedback) and RF (Retest With Feedback) RTs. Asterisk (*) indicates significant difference; white circles are individual means; shading and error bars are SEM.

However, the RT difference returns in the Retest No Feedback block (Figure 3C; *t*_(105)_ = −2.2276, *p* = 0.0280). The presentation of targets during these three blocks were in the same random format, so the reemergence of an RT effect during the Retest No Feedback block is most likely due to the cost of using a strategy since the only instructional change is to recall how to compensate for each rotation when no feedback is given. The purpose of the Retest No Feedback block was to evoke explicit memory of the task from participants since the absence of any cursor feedback would prevent any immediate learning. However, now the group difference is in the opposite direction. The Random group in the Retest No Feedback block has a significantly lower RT compared to the Blocked group that spikes at the beginning and stays relatively higher (Figure 3A). This indicates that the Random group had an easier time than the Blocked group flexibly recalling each strategy they used beforehand. In other words, they were more adept at switching from one strategy to another on a trial-by-trial basis, even though the order was completely unpredictable. However, this significant group effect is not reproduced in the Retest With Feedback block, although it is in the expected direction (Figure 3C; *t*_(105)_ = −1.5703, *p* = 0.1194). Overall, the RT results are in alignment with the contextual interference hypothesis: The Blocked group was faster during training than the Random group and tended to be slower (not significant) during retest.

The second variable we looked at was hand angle. During training and retest, this would correspond to accuracy, and participants were expected to use strategies that included a large explicit component to offset the rotations. In contrast, during the aftereffect and washout blocks, hand angles served as a measure of implicit adaptation. Contextual interference would predict the Random group to outperform the Blocked group and display more compensatory hand angles for all rotations during retest to show they are better able to recall their strategies from training.

The behavioral responses for targets with counterclockwise rotations were flipped so that the data could be averaged as if every rotation was in the clockwise direction (Figure 4A). During training, the Blocked group learns each rotation faster than the Random group (early training [first 10 reaches to every target]: lme: *t*_(104)_ = −2.8363, *p* < 0.01), but there is little difference in hand angle by the end of the block (Figure 4A, B; late training [last 10 reaches to every target]: lme: *t*_(104)_ = −1.0575, *p* = 0.2927). In comparison to Random participants, the Blocked group was able to find and utilize their strategies more quickly, but by the end of the training period, all participants generally exhibited similar reaching behavior. This is all consistent with contextual interference, and the equal performance by the end is most likely due to a ceiling effect.

**Figure 4:**
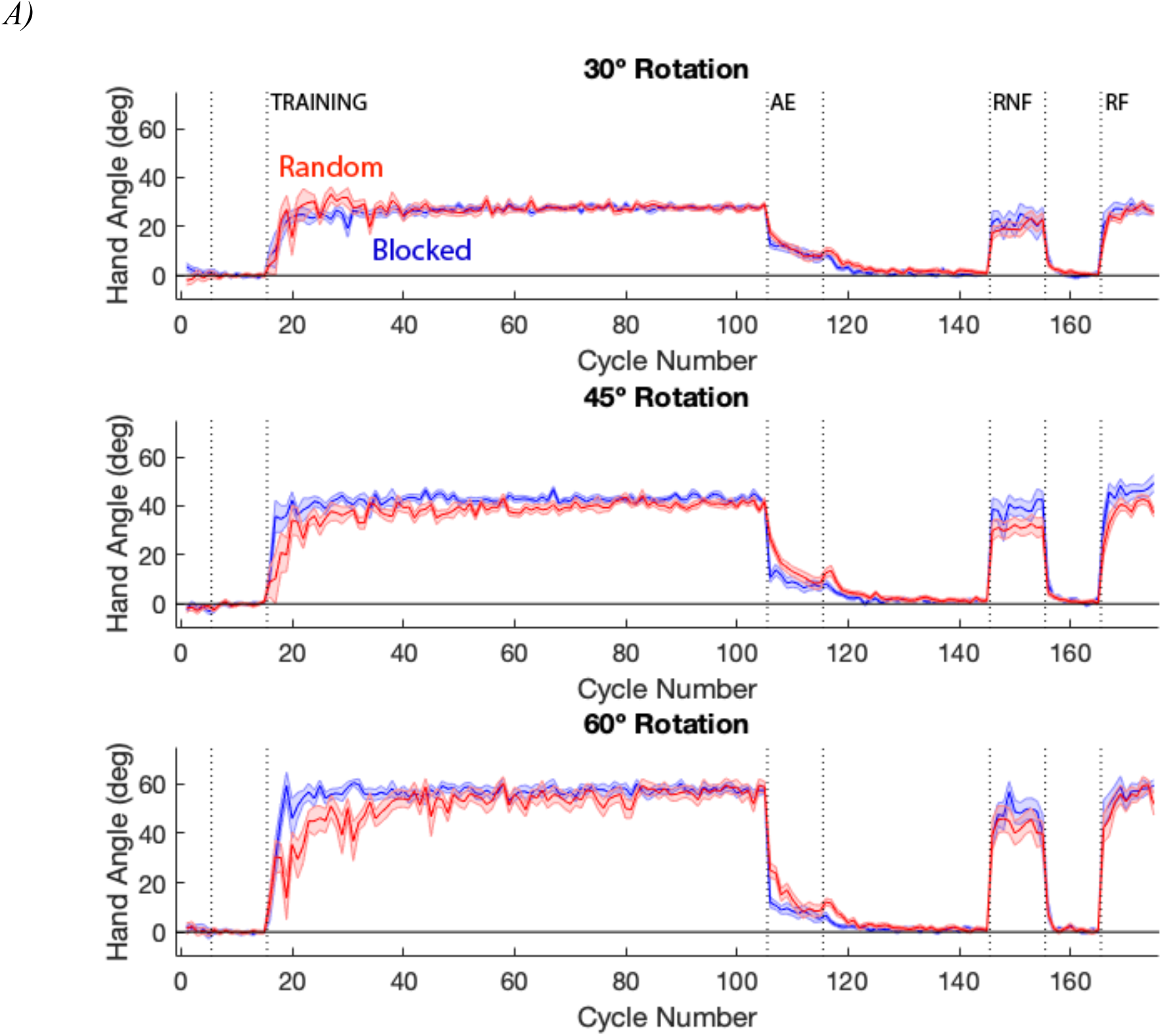
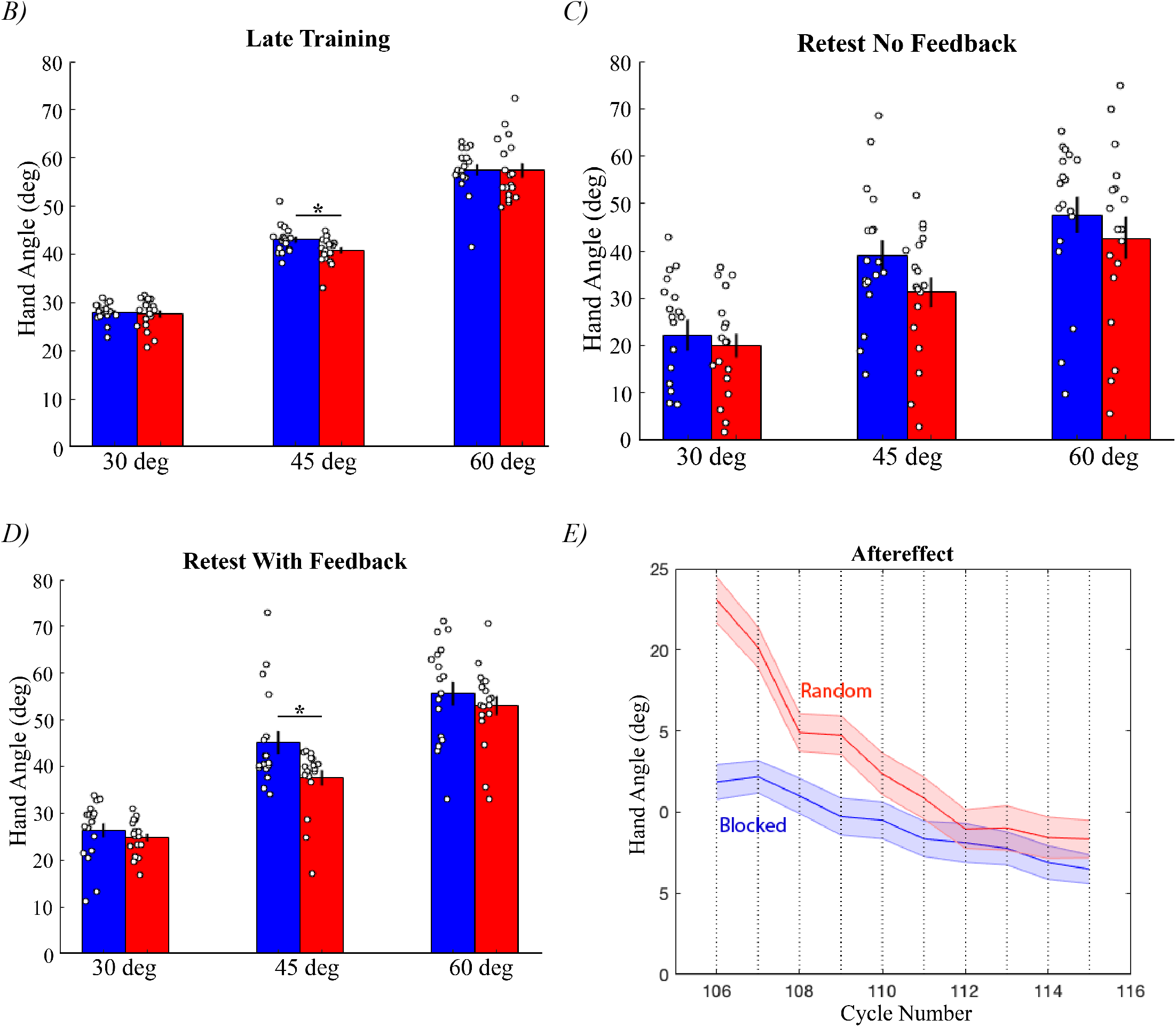
Hand Angles for Experiment 1. **A)** Averaged group endpoint hand angle across Experiment 1 split by rotation. Training, Aftereffect (AE), Retest No Feedback (RNF), and Retest With Feedback (RF) blocks labeled. Data for **B)** late Training (last 10 reaches to each target), **C)** Retest No Feedback, and **D)** Retest With Feedback blocks shown in bar graphs. **E)** Averaged aftereffects for each group for the entire block. Asterisk (*) indicates significant difference; white circles are individual means; shading and error bars are SEM.

The purpose of the Retest No Feedback block was to look at how much explicit memory was retained and then how much of the overall task could be recalled in the Retest With Feedback block. Group hand angles were similar for the retest blocks, making it inconsistent with the contextual interference hypothesis. There was no significant difference between the groups during Retest No Feedback (Figure 4C; lme: *t*_(104)_ = −1.3713, *p* = 0.1732), but there was for the Retest With Feedback block (Figure 4D; lme: *t*_(104)_ = −2.4006, *p* = 0.0181). However, the direction of significance is not in the expected direction; the Blocked group exhibited more compensatory behavior with average reaches closer to the rotation of the cursor. The cost in performance for the Random group seems to return when feedback is re-introduced, indicating that those participants have trouble enacting the correct hand movement and adjusting for their errors. Overall, we do not see any benefit of random practice on the measure of hand angle.

However, we did unexpectedly find a group effect when we assessed implicit adaptation in the Aftereffect block. During this phase of the experiment, there was no visual feedback and the participants were instructed to reach directly towards the target. While both groups showed adaptation, the Random group exhibited a significantly larger aftereffect (Figure 4E; *t*_(105)_ = 2.1667, *p* = 0.0325). Even though the difference slowly diminishes over time, its presence is surprising. From previous studies, we would have predicted no difference in adaptation since it is traditionally thought of as a robust process that would not be influenced by factors such as cursor rotation or target arrangement (Morehead et al., 2017; Kim et al., 2018). All participants experienced the same number of reaches and had the same amount of practice to each target during training. The only point that the groups differ is whether the order of targets was random or blocked, which inexplicably led to different aftereffects. To further examine how and why contextual interference affects implicit adaptation, we conducted a second experiment that replaced the visuomotor rotation with an error-clamp.

### Experiment 2

Task-irrelevant visual feedback has been used to study implicit adaptation without any influence from explicit learning (Shmuelof et al., 2012; Vaswani et al., 2015; Morehead et al., 2017; Kim et al., 2018). For this manipulation, the cursor is fixed to an invariant path that is rotated from the target and independent of the participant’s hand position. Even though participants are fully informed of the manipulation and told to reach their hand directly to the target while ignoring the cursor, they still implicitly adapt and exhibit aftereffects. The task-irrelevance of the cursor perturbation solely induces a sensory prediction error, which is the difference between where the cursor is expected to go (usually in parallel with the hand) and where it actually goes. This error is commonly thought to be what drives implicit adaptation (Morehead et al., 2017).

In Experiment 2, we used this method to ask if random and blocked practice schedules differentially affect implicit adaptation. During training and retest blocks, we provided task-irrelevant clamped visual feedback that was rotated in a similar manner as Experiment 1 (Figure 1B). Participants were told that their goal was to hit the target with their hand and ignore the cursor feedback. Instructions during the retest block was to continue moving to the target since there was no strategy component of the task to remember. Everything else was identical to the previous experiment (Figure 1D).

The difference in RT between the EC-Blocked and EC-Random groups was significant for all rotations throughout the training blocks (Figure 5A, B; late training [last 10 reaches to every target]: *t*_(34)_ = 3.6316, *p* < 0.0001 for 30° rotation; *t*_(34)_ = 3.6105, *p* < 0.0001 for 45° rotation; *t*_(34)_ = 2.1580, *p* = 0.0381 for 60° rotation). Participants in the random condition consistently had longer RTs compared to those in the blocked condition. There was no similar group effect in the Retest No Feedback block (Figure 5A; Retest No Feedback: *t*_(105)_ = 0.0104, *p* = 0.9917), which was expected since participants were not required to explicitly recall any reaching strategies.

**Figure 5:**
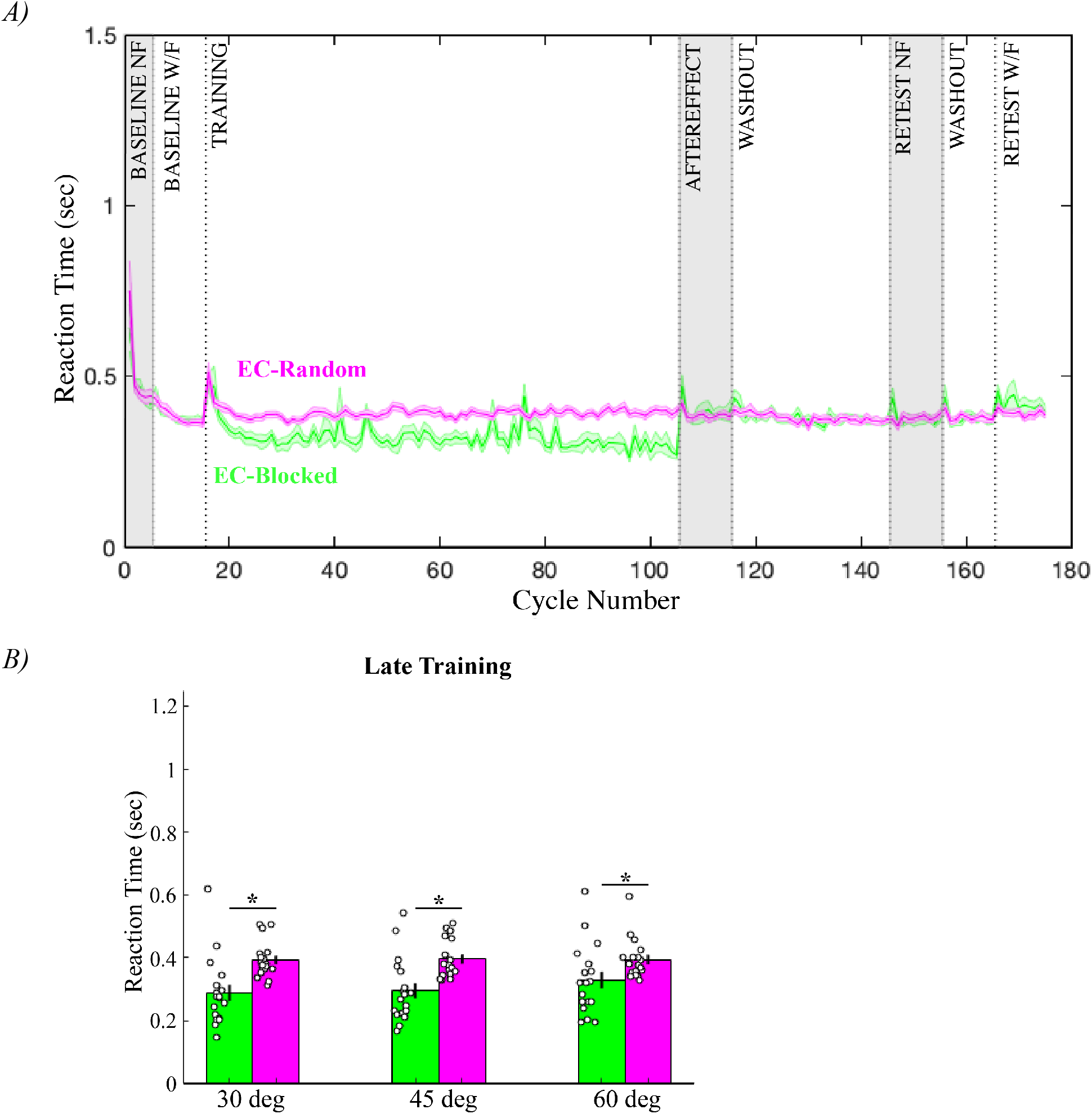
Reaction Time for Experiment 2. **A)** Averaged group RT across Experiment 2 **B)** Late training (last 10 reaches to each target) RTs split by rotation. Asterisk (*) indicates significant difference; white circles are individual means; shading and error bars are SEM.

The rate of adaptation was similar among participants in the EC-Blocked and EC-Random groups for all of the rotations (Figure 6A; entire training: *t*_(34)_ = 0.0786, *p* = 0.9378 for 30° rotation; *t*_(34)_ = 0.9097, *p* < 0.3694 for 45° rotation; *t*_(34)_ = −0.1059, *p* = 0.9162 for 60° rotation). However, there were still significantly different aftereffects after the training period (Figure 6B; *t*_(105)_ = 2.3436, *p* = 0.0210). This suggests that various levels of retention may be playing a role here, which we will further explore below. Hand angles for the retest blocks did not differ between groups, which was expected since there was no task-strategy component to recall (Figure 6A; Retest No Feedback: *t*_(105)_ = 1.7106, *p* = 0.0901; Retest With Feedback: *t*_(105)_ = 0.0194, *p* = 0.9846).

**Figure 6:**
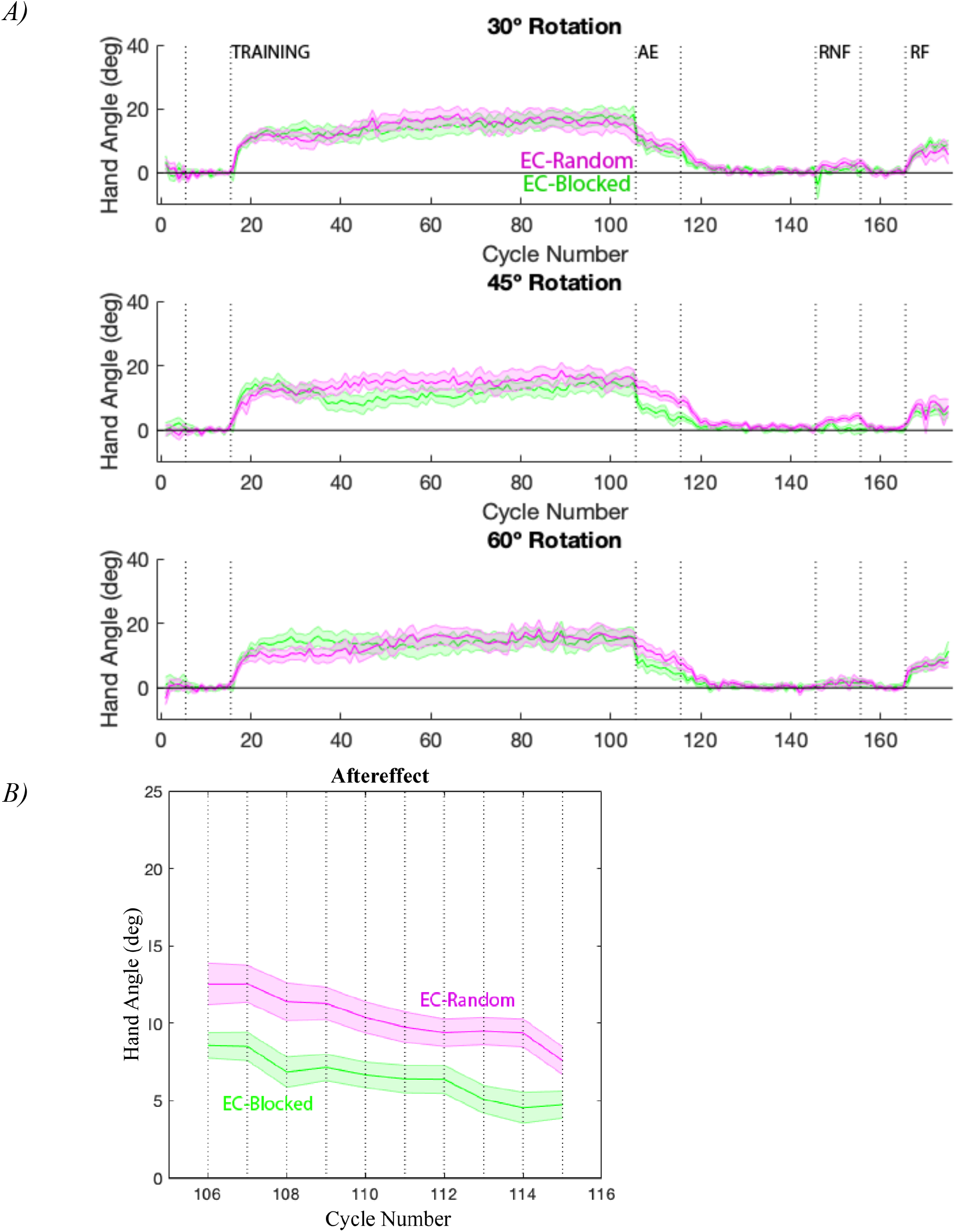
Hand Angles for Experiment 2. **A)** Averaged group endpoint hand angle across Experiment 2 split by rotation. Training, Aftereffect (AE), Retest No Feedback (RNF), and Retest With Feedback (RF) blocks labeled. **B)** Averaged aftereffects for each group for the entire block. Asterisk (*) indicates significant difference; white circles are individual means; shading and error bars are SEM

Altogether, we see an effect of contextual interference on adaptation: Hand angles are similar throughout the training period but then are clearly different during the Aftereffect block, showing a notable influence of target order on implicit learning.

#### Varying Levels of Retention in Implicit Learning

Due to the significant amount of implicit adaptation observed in the Random group from Experiment 1, we performed a second experiment to look at implicit learning and found a similar difference in aftereffects between the groups. To compare across all groups in both experiments, Figure 7A shows the average hand angles for each group during the entire Aftereffect block. The EC-Random group is not significantly different from the Random (*t*_(105)_ = −1.4747, *p* = 0.1433) group. This lack of difference shows that any interaction between implicit and explicit processes that give Random participants a big adaptation boost at the beginning of the block is not significant enough to last until the end. This is similarly seen when the EC-Blocked and Blocked groups are compared (*t*_(105)_ = −1.7917, *p* = 0.0761). Overall, there does not seem to be any significant differences between experiments, which indicates that the experimental difference of using a reaching strategy or ignoring the cursor did not have a great effect. However, the group differences within each experiment (Random vs. Blocked in Experiment 1 and EC-Random vs. EC-Blocked in Experiment 2) suggest that the target order plays some role in influencing how much aftereffect each group has.

**Figure 7:**
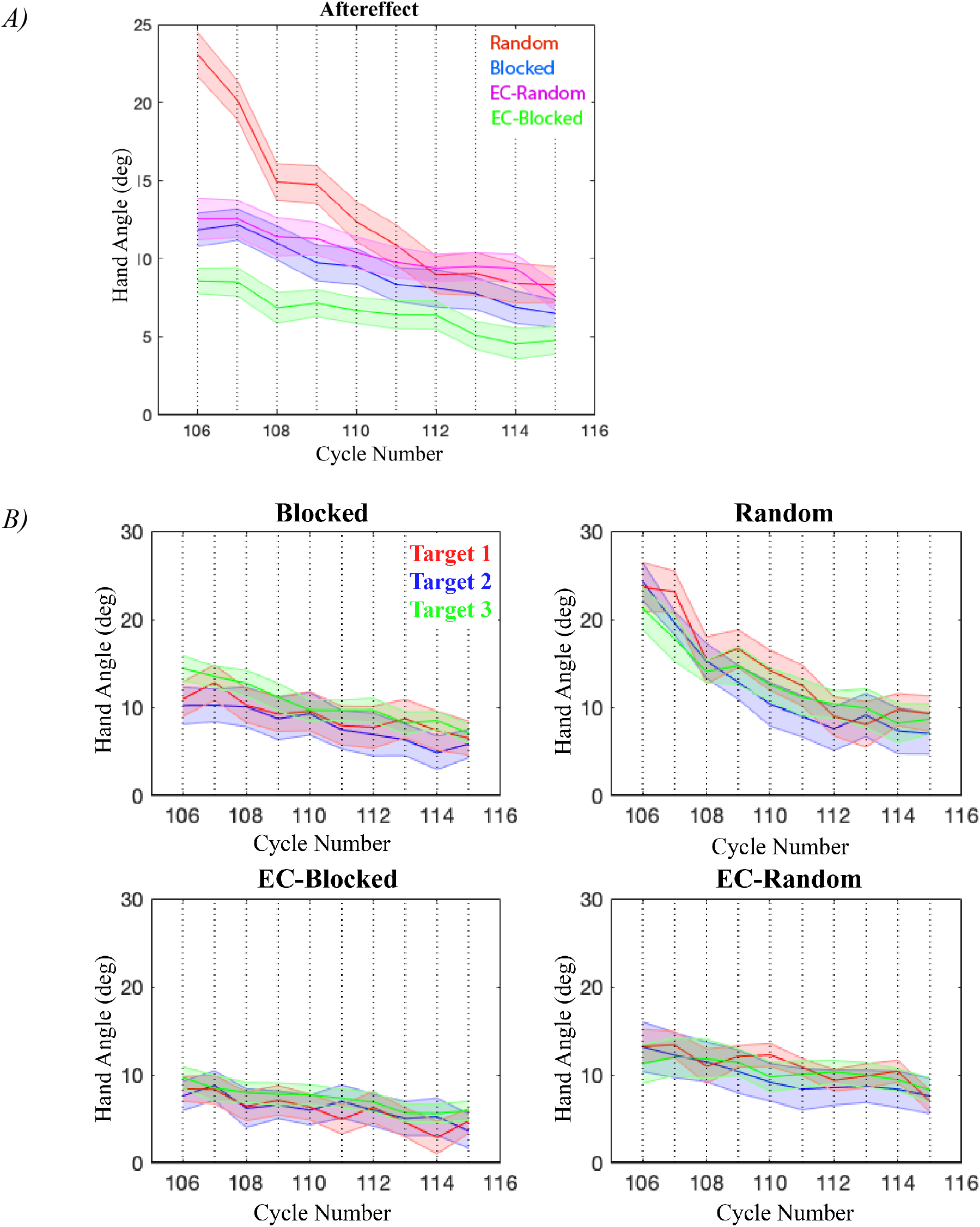
Implicit Adaptation for All Groups. **A)** Averaged aftereffects for all groups across the entire block. **B)** Each group’s aftereffect data split by target. Color code changes: Target 1 (30°) = red, Target 2 (150°) = blue, Target 3 (270°) = green. Shading is SEM.

During training, we observed similar rates of adaptation between error-clamp groups, which shows that implicit adaptation really is an invariant process (Taylor et al., 2014; Morehead et al., 2017; Kim et al., 2018). It is not affected by switching between targets or rotations and, therefore, contextual interference. We split the aftereffect hand angles by target in order to determine whether or not forgetting has to do with the observable difference in aftereffects (Figure 7B). The groups experiencing the blocked training schedule always reached to the 30° target first, then the 150° one, and then 270°. This was done in order to simplify the number of possible configurations since the order of rotations was already being counterbalanced across participants. We used this determine if the recency of the targets’ appearance plays a role. The Random and EC-Random groups served as a control because they reached to all three of the targets right up until the end of the training period, so we reasonably do not see any effect of target order (Figure 7B; Random: *t*_(538)_ = 0.6739, *p* = 0.5006; EC-Random: *t*_(537)_ = 1.1316, *p* = 0.2583). The EC-Blocked group similarly showed no target effect, but the Blocked group did have some significant aftereffects between targets (Figure 7B; Blocked: *t*_(537)_ = 3.8257, *p* < 0.0001; EC-Blocked: *t*_(536)_ = 1.9461, *p* = 0.0522). The fact that the effect is only seen in the Blocked group means that forgetting due to target order plays a bigger role when explicit processes are involved. This points to some interaction between the two motor learning systems. There may be some forgetting involved that gives the EC-Blocked group a lower aftereffect, but it clearly does not tell the whole story. Therefore, there is still some unknown underlying effect that context has on implicit learning.

Altogether, we found that even though contextual interference does not affect the rate of adaptation, it does affect how much is retained and observed during the Aftereffect block. When considered in the light of the results from Experiment 1, random practice seems to provide an advantage in recalling information in the future. Random participants are better able to hold on to the previous strategies they used and are, therefore, faster at choosing which one to perform during retest, just like how they are better able to hold on to any adaptation acquired throughout training. Where they have trouble is using these strategies to reach accurately. This suggests contextual interference provides more of an advantage for maintaining big-picture information rather than the finer details, making it more suited for motor tasks that emphasizes action selection rather than precision.

## Discussion

The purpose of this study was to examine the effects of contextual interference on explicit and implicit motor learning. The results we have collected provide an explanation for why contextual interference is not a global effect that could boost learning in any motor task. We generally observed no difference in retested hand angles between Blocked and Random participants, especially in terms of explicit memory that was specifically tested for in the Retest No Feedback block. We did see a significant group effect in hand angle during the Aftereffect block, which was reproduced in our error-clamp experiment that focused on implicit learning. For the groups in the error-clamp condition, we saw similar rates of adaptation (Training block) but significantly different aftereffects (Aftereffect block), indicating some underlying process of retention that benefits random participants.

We also looked at group RT differences and found participants in both of the random conditions had higher training RTs. There was a 198 ms difference in averaged RT for Random and Blocked groups that dealt with a dual cost of target order and reaching strategy. However, there was only a 71 ms difference between EC-Random and EC-Blocked groups (no reaching strategy), meaning that only 71 ms of the 198 ms difference in Experiment 1 can be attributed to target order alone. In other words, the cost of not knowing which of the three targets would show up is 71 ms. The rest could be due to the time it takes to choose which strategy to use. For the Retest No Feedback block, only the Random group (from Experiment 1) had significantly lower RTs, which indicates that experience with interference in the process of formulating an explicit reaching strategy could help to improve recall later on. We, therefore, believe that contextual interference helps to retain information for longer periods of time that can more easily be accessed later on; however, there is no guarantee that the information is completely accurate. This shows that experiencing interference is more beneficial for motor tasks where the primary goal is selecting an action plan as fast possible.

Our observed benefit of contextual interference to RT could explain why this effect is consistently seen in multi-segment movement tasks popularized by Shea and Morgan (1979). Participants in these experiments are usually given three different sequences to learn and remember that are each associated with their own stimulus. For example, a green light will signal one sequence of movements to be performed as fast as possible, while a red light will signal another. Those in the random practice condition were able to perform each sequence significantly faster when they are later retested, showing an enhanced ability to recall which movement to do when the appropriate stimulus appears. This was similarly seen in our experiment. When the Retest No Feedback block started, Random participants were more adept at flexibly switching between strategies than Blocked participants. Contextual interference seems to help by strengthening the association of the correct movement to its stimulus. This does provide support to Battig’s initial hypothesis that the effect of contextual interference is due to a higher engagement of mental processes.

Similarly, the lack of any noticeable effect in hand angle could explain why interference during practice rarely led to any performance benefits in non-laboratory experiments. One study by Bortoli et al (1992) focused on applying contextual interference to learning different volleyball skills. Their experiment consisted of learning a bump, two-hand volley, and an under-hand serve in order to propel the volleyball across the net and hit a target drawn on the floor. Participants were awarded points during practice (with or without interference) based on how accurately they could hit the center of that target, which were used to measure performance. After a long training period, they were retested to assess how much learning was retained. Overall, Bortoli et al (1992) found no significant difference in retest scores among the differently scheduled practice groups; those in the random condition did not show any benefit in performance, so contextual interference was not helpful in this motor learning task. In their discussion, they considered that one of the reasons behind this lack of effect could be due to the strong emphasis they placed on accuracy (getting the ball to hit as close to the target center as possible to maximize the score), which we find support for with our own hand angle results. Additionally, another experiment had the same task design but instead used basketball shots from different positions on the court and scored how many successful baskets were made (Landin & Hebert, 1997). They also found no benefits of random practice on learning. We believe that placing an importance on hitting a target with a cursor, a volleyball, or a basketball effectively makes this type of motor task resistant to the benefits of contextual interference. One possible reason may be because achieving and then retaining accuracy may require immediate corrections that a random practice schedule does not allow. Further research will be needed, but we are confident that interference during training generally does not improve the accuracy of a movement.

A new finding in our study is that contextual interference influences the amount of adaptation retained in the random groups. In support of the current model of adaptation, we did observe the rate of adaptation to be comparable between EC-Blocked and EC-Random groups, consistent with the hypothesis that this implicit process is invariant and automatic (Morehead et al., 2017; Taylor et al., 2014). However, we still saw significantly different aftereffects between Blocked and Random groups and between EC-Blocked and EC-Random groups. We initially attributed this effect to some forgetting, which we parsed out by separating hand angles during the Aftereffect block by target (Figure 7B). This will be especially revealing for the blocked groups who reached to the 30° target two blocks ago and the 150° target one block ago. We only found a significant target effect for the Blocked group and not the EC-Blocked group, which means forgetting may play a role but does not fully explain why blocked participants show less adaptation. Through an unknown process that needs more exploring, interference somehow enables random participants to retain for a longer amount of time any adaptation they acquired through training. The Blocked group does seem to be the most impacted, which may mean that the interaction of forgetting and interference on implicit learning could be exacerbated by the addition of explicit processes. For example, any adaptation acquired from reaching to the 30° target may have been lost faster in the Blocked group than the EC-Blocked group due to the simultaneous use of explicit and implicit processes attempting to compensate for the rotation at the 150° target. By the time Blocked participants begin the Aftereffect block, the amount of adaptation for the 30° and 150° targets have been significantly reduced, pulling down the group average and giving the observable group effect. The impact of explicit processes on adaptation is similarly observed when comparing Random and EC-Random groups. There is some interaction between these two learning components that previous studies have touched on that needs to be further examined (Taylor et al., 2014; Bond & Taylor, 2015).

There are a few limitations to our study. The first is that our experiment was not specifically designed to look at RT. If it was, we could have included an experimental component that more closely looked at a trade-off between RT and accuracy. For example, after the target appears, a set amount of time all participants have to wait before they can start reaching. By varying the wait time, we could see how random and blocked participants changed their behavior and how much more they actually benefit from having more time. Our RT results are indicative of a significant contextual interference effect that a more specific RT-designed experiment could explore.

The second is that our overall task was easy enough that participants were able to quickly relearn it in the retest blocks. We could have increased the cognitive load by introducing more targets and more rotations and then see clear observable effects of contextual interference.

In summary, our utilization of a simple reaching task showed contextual interference effects in RT and adaptation but not in explicit. We believe this means that experiencing interference during training boosts participants’ ability to retain information that helps them exhibit increased aftereffects or better recall which strategy to choose but does not help with accuracy. Contextual interference is the more beneficial for situations that require fast recall, which has implications for how this effect can be applied to motor learning in the real world.

